# The Impact of Peri-implantitis on the Proteome Biology of Crevicular Fluid: A pilot study

**DOI:** 10.1101/2022.12.01.518583

**Authors:** Tim Halstenbach, Katja Nelson, Gerhard Iglhaut, Oliver Schilling, Tobias Fretwurst

## Abstract

**Background:** The proteome of the peri-implant crevicular fluid (PICF) has not been systematically investigated. The aim of the present study was to reveal the proteome biology of dental implants affected with peri-implantitis.

**Methods:** Patients with at least one diseased implant were included (probing depth ≥ 6 mm, ≥ 3 mm peri-implant radiological bone loss). Using sterile paper strips, samples were collected from healthy implants (I), healthy teeth (T) and peri-implantitis affected implants (P). Proteome analysis was performed using liquid chromatography – tandem mass spectrometry (LC-MS/MS) and data independent acquisition, allowing the identification and quantification of human and bacterial proteins as well as semi-specific peptides.

**Results:** 38 samples from 13 patients were included in the study. 2332 different human proteins were identified across all samples. No differentially expressed proteins between T and I were found. Comparing P to I, 59 proteins were found upregulated and 31 downregulated in P with significance. Upregulated proteins included proinflammatory proteins such as immunoglobulins, dysferlin and S100P, as well as antimicrobial proteins, e.g. myeloperoxidase or azurocidin. Gene ontology analysis further revealed higher activity of immunological pathways. Proteolytic patterns indicated the activity of inflammatory proteins such as cathepsin G. 334 bacterial proteins were identified and quantified. Peri-implantitis showed elevated proteolytic activity.

**Conclusion:** I and T share similarities in their proteome, while diseased implants deviate strongly from healthy conditions. The PICF proteome of peri-implantitis affected sites exhibits an inflammatory fingerprint, dominated by neutrophile activity when compared to healthy implants.

**Summary:** Proteomic analysis of the peri-implant crevicular fluid revealed distinct proteome alterations in peri-implantitis when compared to healthy implants and teeth, while healthy teeth and implants share strong similarities.

## Introduction

Long-term survival rates of dental implants are documented with over 90% after ten years ^12^. However, peri-implant inflammation (peri-implant mucositis: PM; peri-implantitis: P) is common with a prevalence of up to 20% for peri-implantitis and up to 40% for peri-implant mucositis ^3^. Equivalent to periodontology, the etiology was assumed to be determined by a microbial biofilm in a susceptible host. Prosthetic, surgical or biomechanical factors are discussed as an additional etiological mechanism for peri-implant bone loss ^4–6^. Currently, no standardized therapy of diseased implants is available, the treatment options are associated with high recurrence rates even after a decade of intensive research ^7,8^. Drug therapies with a curative approach are not available ^9^.

Early non-invasive diagnostic tools or parameters are missing, only invasive, unspecific diagnostic tools e.g. peri-implant pocket depth measurement, radiological peri-implant bone level are available to detect peri-implantitis ^10^. Radiological bone loss and probing depth indicate peri-implantitis when the disease has already progressed. The microbiological analysis of peri-implant disease has not revealed any consistent, disease-specific microbiome profiles and can therefore not be used for diagnosis or disease monitoring ^11,12^. In the field of periodontology and implantology, potential diagnostic markers of periodontitis in the gingival crevicular fluid (GCF) and for peri-implantitis in the peri-implant sulcus fluid (PICF) are discussed as a point-of-care testing laboratory diagnostics ^13^. Potential diagnostic, prognostic or therapeutic markers include enzymes, inflammation and host reaction mediators e.g. cytokines, interleukins as well as metabolic markers related to bone tissue degeneration around the tooth or the implant. For the detection of the markers targeted methods e.g. ELISA, spectrophotometry or flow cytometry were mainly used ^14–17^. In the targeted approach potential diagnostic markers must be selected a priori, and only a limited number of markers can be evaluated. Due to the limitation of the analytical methods used only a small number of proteins have been evaluated, the proteomic composition of the sulcus fluid has not been evaluated to date.

Immunohistological examination of inflamed peri-implant und periodontal tissues demonstrated larger areas of inflammatory infiltrates in peri-implantitis compared to periodontitis. Furthermore, macrophages (CD-68 positive cells), plasma cells (CD-138 positive cells) and neutrophils (myeloperoxidase (MPO) -positive cells) are detected in the peri-implant infiltrates ^18^. Inflamed peri-implant tissue demonstrates a pro-inflammatory M1 macrophage polarization and a lymphocyte-dominated inflammation with interindividual, patient-specific differences. The concept of patient-specific immune profiles in peri-implantitis was corroborated by transcriptome sequencing of the peri-implant inflammatory tissue and has been proposed to be regarded for risk stratification for the therapeutic success of surgical peri-implant therapy ^19^.

In personalized cancer precision medicine, molecular information/profiling of tumors, the microenvironment and the diseased individual is increasingly used for diagnostics and treatment decisions. Mass spectrometry (MS) based proteomics enables quantification and identification of almost all proteins in the medium allowing for multibiomarker approaches and non-targeted exploratory essays ^20^. In the field of periodontology, targeted proteomic methods enabled the evaluation of approximately 100 proteins ^21^, when using MS based proteomics more than 600 proteins were identified in the GCF of periodontitis affected teeth. Studies demonstrated increased expressions of apolipoproteins, immunoglobulins as well as cytoskeletal proteins and histones of neutrophil extracellular traps (NETs) ^22,23^. The PICF proteome has not been investigated by comparing healthy and diseased implants nor has the proteomic composition found in PICF been compared to the composition of GCF. MS-based proteomics of PICF has only been performed once, not including healthy implants or healthy teeth as a control group ^24^, therefore not allowing for the identification of disease specific markers or proteomic clusters.

The objective of the present study is to analyze and compare the proteomic composition of GCF of healthy teeth and PICF of healthy implants and implants with peri-implantitis in order to gain new insights into the disease biology. The hypothesis is that healthy implants and healthy teeth have a different specific sulcus protein signature than diseased implants with peri-implantitis. Furthermore, MS-based proteomics may enable identification of bacterial proteins in the PICF.

## Material and Methods

The study was approved by the ethics committee of the University Medical Center Freiburg, Germany (Ethik-Kommission Albert-Ludwigs-Universität, Freiburg, Votum 337/04). Before enrollment, the patients received information regarding the purpose of the study and signed an informed consent. All patients were consecutively enrolled between 2020 and 2021 (Department of Oral- and Craniomaxillofacial Surgery / Translational Implantology, University Medical Center Freiburg). The study was performed in accordance with the Helsinki Declaration of 1964, as revised in 2013. The study was conducted in accordance with the SRQR (Standards for Reporting Qualitative Research) guidelines.

### Study cohort

Patients with at least one diseased implant were included. Peri-implantitis was defined according to the current international guideline with peri-implant pocket depth ≥ 6 mm, ≥ 3 mm peri-implant radiological bone loss with bleeding and / or suppuration on probing (Berglundh 2018). Minimum age of the participants was 18 years. Prosthetically restored implants were included which had received the restoration > 12 months prior. Screw retained- and cemented restorations were included.

Patients were excluded from the study if one of the following conditions was given: diabetes, immunosuppression or immunosuppressive medication, current radiation of the head or neck area, or oral mucosal diseases (e.g. erosive lichen planus), bisphosphonate or antiresorptive agent intake, current untreated periodontal disease or gingivitis, pocket depth of ≥ 4 mm at the teeth immediately next to the dental implant to be examined, severe bruxism and patients with inadequate oral hygiene or not motivated to provide adequate oral care at home, current intake of antibiotics or local application of antibiotics. Only patients that had not a received a peri-implantitis treatment shortly before the time point of implant removal were included. All patients were frequently examined by professionals and periodontally treated, if necessary, therefore included patients did not present active periodontitis sites. Hence, in this proof-of-concept study no samples of periodontitis affected teeth were included.

### PICF sample collection

PICF sample collection was performed as described^17^. Briefly, after gentle air-drying, peri-implant sulcus fluid samples were obtained the target implant site, as well as at the healthy tooth and healthy implant, with sterile paper strips^*^. The strips were placed into the sulcus with a minimum depth of 1–2 mm for 30 sec. The procedure was performed three times each time using a new strip. Strips were instantly collected in cryotubes and frozen using liquid nitrogen. Subsequently, samples were transported on ice and stored at – 80 °C.

### Sample Preparation for LC–MS/MS

Samples were thawed and eluted in 150 μl of 0,1 % acide labile surfactant in 0.1 M HEPES. After sonification for 15 min, samples were incubated at 90 °C for 30 min. Reduction was performed by adding a final concentration of 5 mM DTT and incubating for 30 min at 37 °C, followed by alkylation in 15 mM chloracetamide and again incubating for 30 min at 37 °C in the dark ^25^. For tryptic digestion 1 μg LysC^†^ was added and incubated for 2 hours at 37 °C. Further, 1 μg of sequencing grade trypsin^‡^ was added. Digestion was performed over night at 37 °C. By adding TFA to a final concentration of 2 % the acide labile surfactant was precipitated and after centrifugation for 15 minutes at 15000 rpm (21130 rcf) the supernatant was transferred to iST-collumns^§^. Desalting was performed following manufacturer’s instructions. In short, columns were washed four times with two kinds of washing buffers (washing-buffer I: 1 % TFA in 2-propanol; washing-buffer II: 0,2 % TFA in ddH_2_O). Peptides were eluted into low bind Eppendorf-tubes with 200 μl of 2 % triethylamine in 80 % acetonitrile (in water). Peptide amount was measured with a bicinchoninic acid assay. To generate a spectral library, 5 μg of each sample with more than 10 μg of peptides were pooled. Until MS, samples were dried and stored at – 80 °C.

### MS Measurements

All samples were analyzed on an Orbitrap mass spectrometer^**^ coupled to an Easy nanoLCTM 100 UHPLC^††^. 0.5 μg of peptides were injected and iRT peptides^‡‡^ were added before MS measurements. Analytical columns were self-packed with C18-coated silica beads (Reprosil Pur C18-AQ, d = 3 Â)^§§^. Buffer A was 0.1 % formic acid, buffer B was 0.1 % formic acid in 80 % Acetonitrile.

To generate an extensive spectral library, the pooled sample was injected for MS six consecutive times with different mass ranges (mass ranges: 400-500, 500-600, 600-700, 700-800, 800-900, 900-1000 Th). A two-step gradient (110 min) was applied with Buffer B increasing from 8 % to 43 % over 90 min and from 43 to 65 % in 20 min. The mass spectrometer was set to Data Independent Acquisition (DIA)-mode with the following settings: AGC target: 1e6; maximum injection time: 80ms; NCE: 25 and 30; MS1 resolution: 70,000; MS2 resolution: 17,500.

Samples were measured on the same column in DIA-mode. Mass-to-charge range was 400 – 1000 m/z, two cycles of 24 m/z widows were applied with a 50 % shift inbetween cycles.

### Generation of Spectral Library and Sample Analysis

Processing of RAW-files from MS-measurements was performed using DIA-NN (v. 1.7.12) ^26^. DIA-NN is a neural networks-based application for deeper proteome decryption of MS-data. An UNIPROT human database was used, (downloaded in June, 2021) in combination with iRT protein sequences^***^ to create a spectral library. In DIA-NN, tryptic digestion was chosen, allowing for one missed cleavage site. Minimum peptide length was set to seven. Analysis allowed for cysteine carbamidomethylation. Other modifications were not included. Precursor charge range was set to 1 - 4 and precursor m/z range was 300-1800 Th. Fragment ion m/z range was set to 200 – 1800 Th. Precursor false discovery rate was 0.01. The spectral library was subsequently used for sample analysis.

Same settings regarding FDR, peptide length and peptide modification were used for analysis of individual samples. In addition, match between runs was enabled. All samples were analyzed in the same run. The protein groups (PG) normalized output of DIA-NN was used for further analysis

### Identification of bacterial proteins

For identification of bacterial proteins, a second DIA-NN analysis was performed. Genomes of bacteria found in the oral cavity were downloaded from the NIH Human Microbiome Project Reference Genome Sequence Database in September 2021. The reviewed human database that was used for the detection of human proteins was combined with the microbiome genomes. The fasta-file compiling both human and bacterial proteins was imported into DIA-NN for spectral library generation. Otherwise, the same settings as listed above applied for library generation and subsequent sample analysis. For further analysis, only proteotyptic peptides were allowed.

### Identification of Semi-specific Peptides

In a third DIA-NN analysis, *in silico* library generation with DIA-NN based on a reviewed human fasta-database (downloaded from Uniprot in November 2020) was performed. In short, library free search was enabled with a fragment ion m/z-range between 200-1800 Th. N-terminal methionine excision was enabled, *in silico* digestion with cuts at K and R and no missed cleavages were allowed. For analysis of samples, the same settings as mentioned above applied.

For identification of semi-specific and non-tryptic peptides, all peptides found were searched against the fasta-file that had been generated previously in DIA-NN.

### Statistical Analysis

For statistical analysis, RStudio was used (v. 1.4.1103-4) (R Foundation for Statistical Computing). Contaminants, iRTs and reverse entries were removed. For further analysis, only those proteins found in 20 % or more samples were considered. M/Z-values were log2-transformed and median centered. Partial Least Squares Regression Analysis (PLS-DA) and Linear Models for Microarray Data (LIMMA) ^27^ were applied, comparing all three groups (teeth, healthy implant, peri-implantitis), as well as subgroups healthy tooth vs. healthy implant and healthy implant vs. peri-implantitis. The adjusted p-value was set to p_adj_ = 0.05 and only proteins with log2-changes of +-50 % were considered as up- or downregulated. Subsequently a two one-sided test (*TOST*) was chosen for statistical equivalence testing between healthy condition as well as healthy and diseased implants. A minimum difference in logarithmic fold change (logFC) of 1 was chosen, the adjusted p-value was set to p_adj_ = 0.05.

To evaluate how frequently bacterial proteins were identified per condition, missingness of proteins was calculated. Missingness represents the proportion of samples of one condition in which a given protein was not identified. High missingness in one condition indicates that the protein was identified only in a small number of samples.

## Results

38 samples of 14 patients (6 female, 8 male, age range 52 – 85 years, median age was 70.7) were included in the study (Supplementary Table 1). Patients were diagnosed with peri-implantitis according to the above-mentioned criteria. Diseased implants were included and if present a healthy implant and/or healthy tooth in the same patient: six patients with samples of peri-implantitis affected implants (P), healthy implants (I) and healthy teeth (T), three patients with P and I, three patients with P and T and 2 patients with no other sample than P were included. At total, 17 P-, 12 I- and 9 T-samples were included.

### Human Proteome Coverage

2332 different human proteins were identified across all samples. Protein identifications per sample ranged from 724 to 1804 with a false discovery rate < 1%. Average proteome coverage per condition revealed significantly lower protein counts for the P-subgroup when compared to healthy implants (Kruskal-Wallis and subsequent Dunns post-hoc test (p_adj_ = 0.021)) (Figure 1A). No significant differences in protein coverage were found between P and I or between healthy conditions (T and I). We identified a coreset of 357 proteins found in all samples. For further analysis, we only considered proteins found and quantified in at least 80% of all samples of one subgroup (max. 3 missing values in subgroup P, max. 2 missing values in subgroup I, max. 1 missing value in subgroup T). For all three conditions 703 proteins were found in 80% of all samples, for P vs. I 810 proteins and for I vs. T 942 proteins.

**Figure 1.**
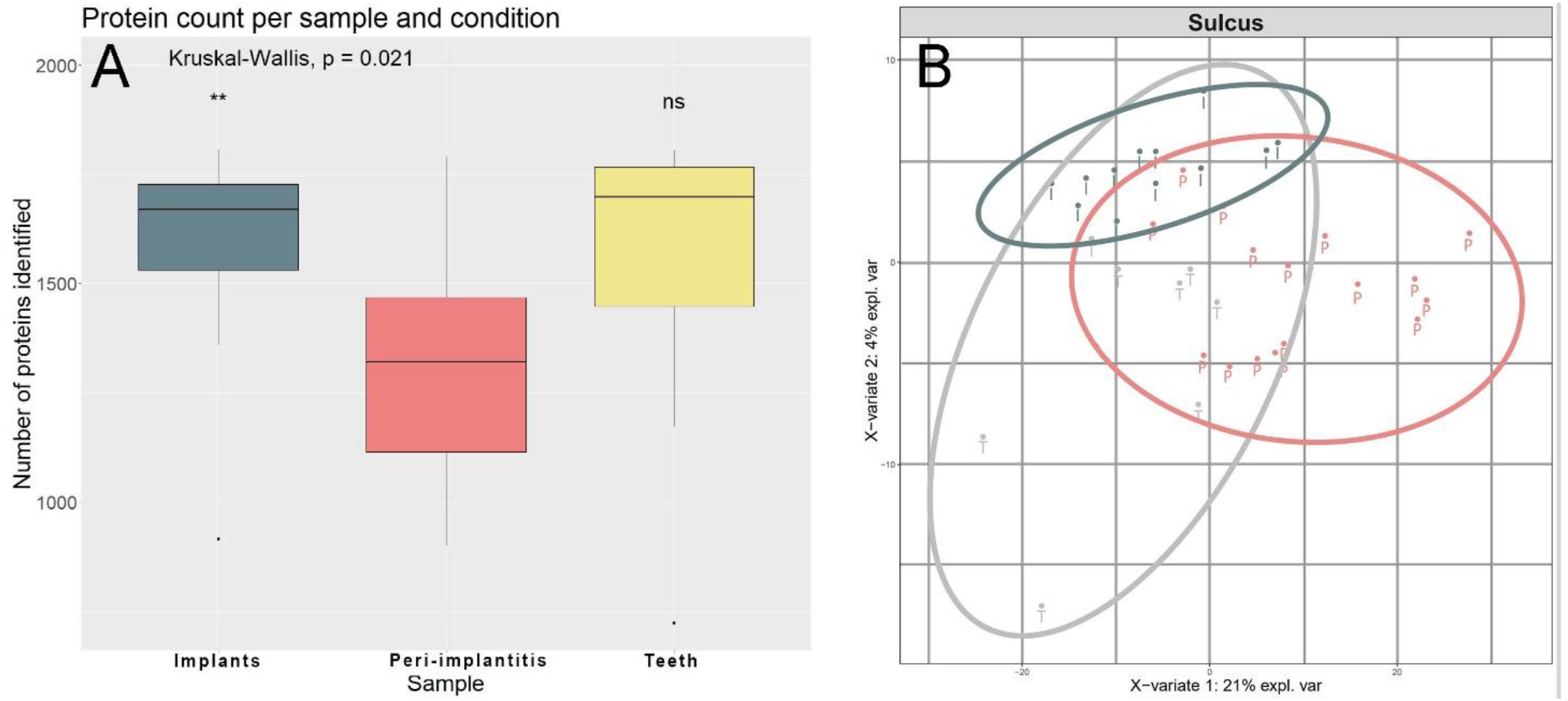
**A**: Proteome coverage for different conditions. Kruskal-Wallis and subsequent Dunns post-hoc tests revealed significant differences between diseased and healthy implants but not between peri-implantitis and healthy teeth. **B**: Partial Least Square Regression Analysis (PLS-DA) of all samples demonstrated a separation of peri-implantitis samples (P) from healthy samples (implants (I) or teeth (T)) and strong similarities between healthy conditions.

### Partial Least Square Regression Analysis and *LIMMA* Indicate Distinct Crevicular Fluid Proteome Profiles for Peri-implantitis as Compared to Healthy Teeth and Healthy Implants

To check for clustering of samples a Partial Least Square Regression Analysis was performed (Figure 1B**Fehler! Verweisquelle konnte nicht gefunden werden**.). PLS-DA resembles a supervised method to distinguish between expected and observed variables. Here, healthy implants were identified as a subgroup of healthy teeth on the proteome level. Peri-implantitis is separated from healthy teeth and healthy implants. This finding underlines a distinguishable proteome profile for crevicular fluid stemming from peri-implantitis as compared to healthy implants or healthy teeth.

For further statistical analysis, *LIMMA* was applied to the following comparisons: P vs. I (Figure 2A); T vs. I (Figure 2B). The latter revealed no significantly expressed proteins, indicating strong similarities between the crevicular fluid of healthy teeth and implants on the proteome level (Figure 2B). When comparing P to healthy implants, 59 proteins were found upregulated and 31 downregulated in P with statistical significance of adjusted p-value ≤ 0.05 (Figure 2A). Here, a variety of inflammatory proteins (dysferlin, immunoglobulin heavy variable 2-70D, leukotriene A-4 hydrolase (LTA4H), S100P and others) and proteins related to bacterial defense (myeloperoxidase, bactericidal permeability-increasing protein, azurocidin and others) were identified.

**Figure 2.**
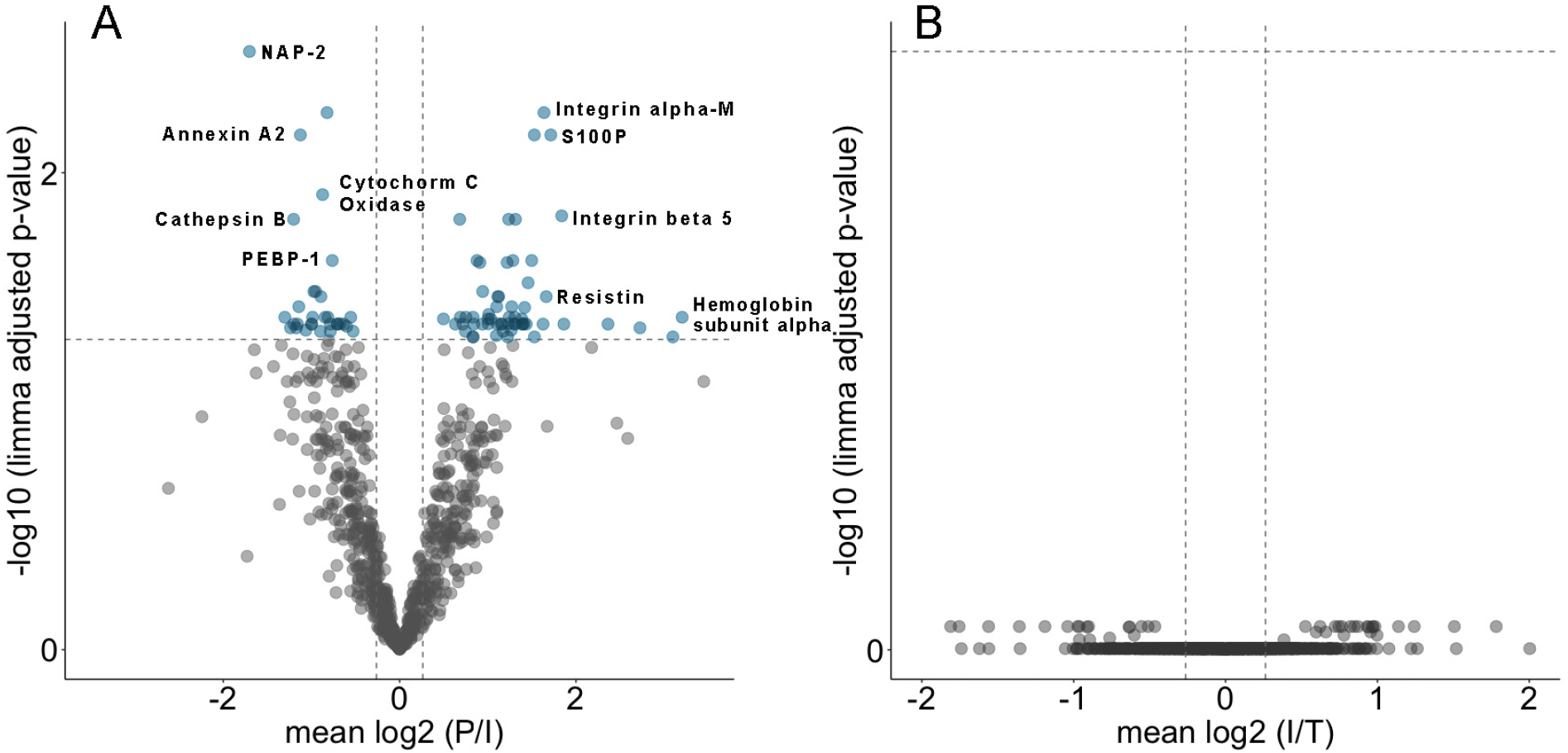
Volcano plot based on *LIMMA* results of peri-implantitis vs. healthy implants (**A**) and healthy implants vs. healthy teeth (**B**). log2 cut off: 50%. Adjusted p-value <=0.05. Colored dots represent up- or down-regulated proteins. A: *LIMMA* demonstrates strong differences between the crevicular of diseased and healthy implants on the proteome level. B: *LIMMA* demonstrates strong similarities between the crevicular of healthy implants and healthy teeth on the proteome level.

To further compare differences and similarities between the conditions on a proteome wide basis, a two one-sided test (*TOST*) was performed subsequent to *LIMMA*. Numbers of proteins identified in equivalent intensities according to TOST and proteins identified with differential expression between conditions P vs. I and I vs. T are depicted in Figure 3.

**Figure 3.**
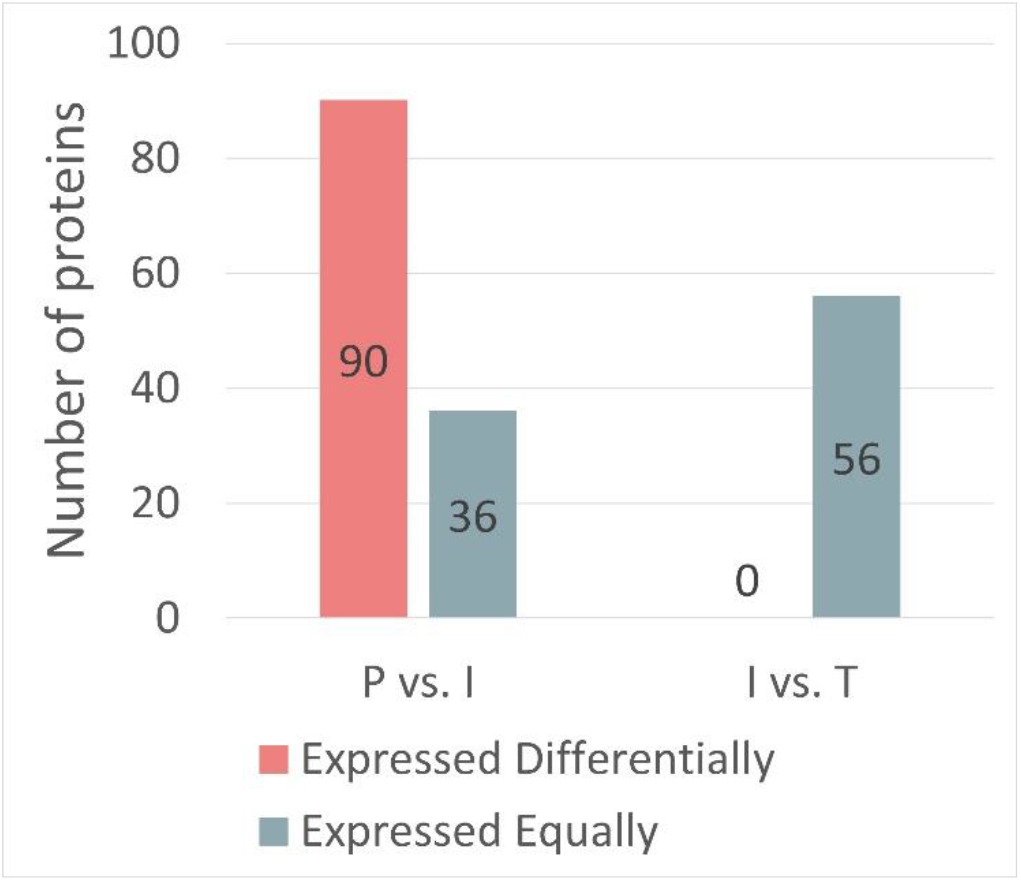
Bar plots of proteins identified as differentially expressed according to LIMMA and proteins expressed equally according to TOST for comparisons P vs. I and I vs. T. Peri-implanitits (P), healthy teeth (T) and healthy implants (I). LIMMA and subsequent TOST demonstrate strong similarities between the healthy conditions and significant differences between healthy and diseased implants.

### Matrix Metalloproteinases-8 and -9

In total, 21 proteases related to inflammation were identified and quantified across all samples. Both Matrix Metalloproteinases (MMP)8 and MMP9 were found in all samples with elevated levels in the P subgroup (Supplementary Table 2). Moderated p-value (P vs. I) for MMP8 was p = 0.037 and p = 0.042 for MMP9 (Supplementary Table 2). Although their adjusted p-values exceeded p_adj_ = 0.05 (p_adj_ < 0.2; P *vs*. I), MMP8 and -9 remain of interest in the context of peri-implantitis since they are endoproteolytic enzymes with collagenolytic / gelatinolytic activity. Leukotriene A-4 hydrolase (LTA4H) and gamma-glutamyl hydrolase were found upregulated in P with statistical significance (p_adj_ ≤ 0.05), yet these enzymes are not considered prototypical endoproteases.

### Gene Ontology Enrichment Analysis

A gene ontology (TopGO) enrichment was performed to categorize proteins identified as differentially expressed. Proteins up- or downregulated around diseased implants when compared to healthy implants were included (log2-fold change of +-50 %, p_mod_ ≤ 0.05). TopGo enrichment revealed several immunological pathways to be upregulated with strongest manifestation in “humoral immune response”, “defense response to other organism” and “regulated exocytosis” (Figure 4). Naturally, inflammatory pathways show great overlap in the associated proteins. For example, both “response to other organism” and “cell killing” include cathepsin G, azurocidin or cathelicidin antimicrobial peptide. In contrast, for I an increase in “muscle structure development” was observed, with multiple structural proteins assigned to the pathway (lamins, desmoplakin, myotrophin and others).

**Figure 4.**
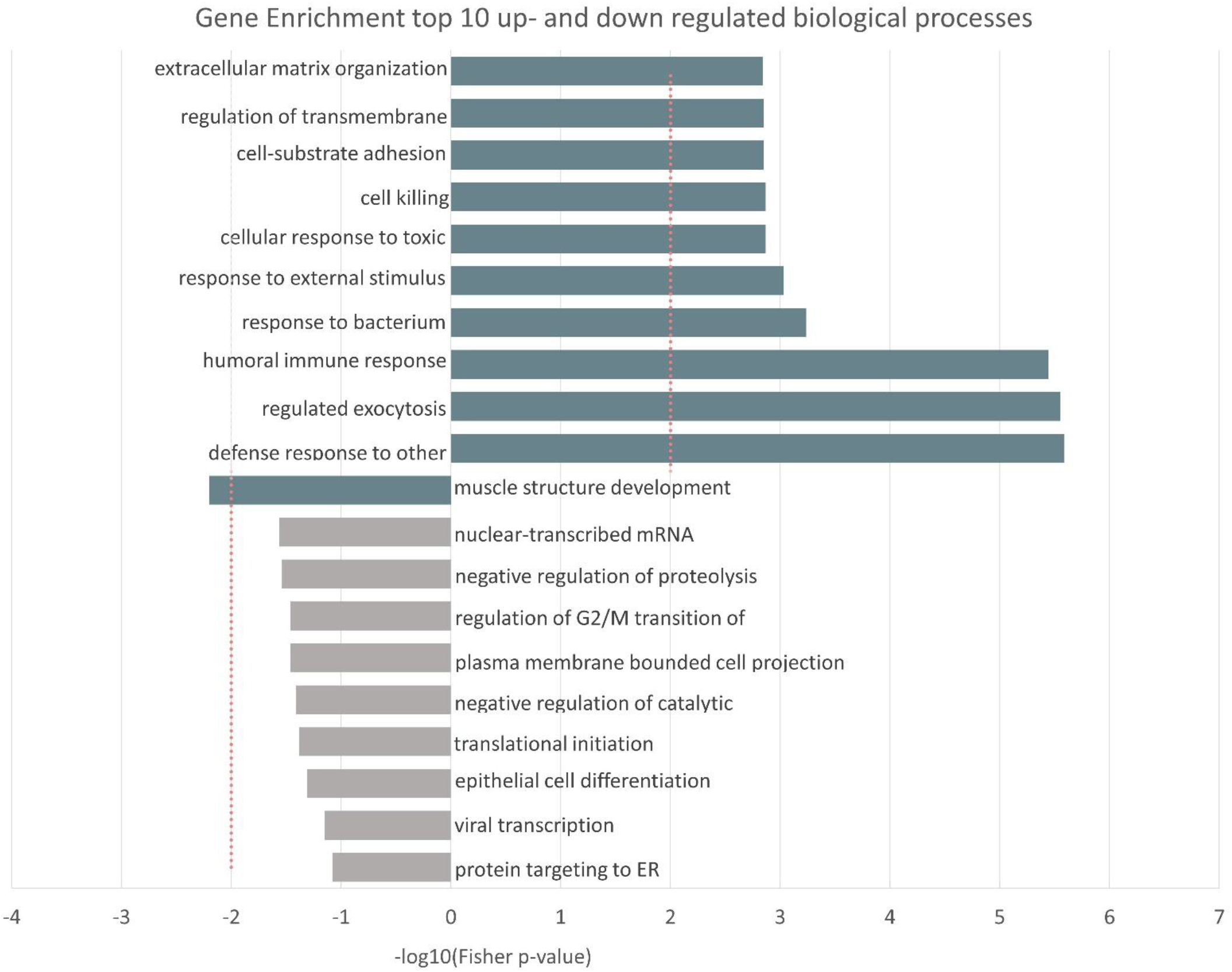
Gene Ontology-analysis of differentially expressed proteins of peri-implantitis (P) vs. healthy implants (I). Top ten up- and downregulated biological processes listed. Upregulated processes: right, downregulated pathways: left. Differentially expressed proteins found in *LIMMA* (p_mod_≤0.05) were used for TopGO-Analysis. Meta-Analysis performed using Fisher’s method. Results interpreted as statistically signficant if -log10(Fisher p-value) > 2.0 or < −2 (dashed lines). TopGo-Analysis demonstrated increased immune response pathways in diseased samples.

### Endogenous Proteolysis

In classical bottom-up proteomics using trypsination, endogenous proteolysis manifests in the form of truncated, so-called semi-specific peptides. To study endogenous proteolysis in peri-implantitis, we performed an additional data analysis step with a spectral library that also represents N- or C-terminally truncated peptides of the human proteome. We note that this approach cannot capture endogenous proteolysis with trypsin-like specificity. 10566 different peptides were identified, 2362 of which were of semi-specific nature. Proportional intensities of semitryptic peptides were calculated per sample (Figure 5). Increased proportional intensities of semi-specific peptides were found in P with statistical significance compared to healthy teeth (p = 0.0057), indicating that peri-implantitis lesions possess higher endogenous proteolytic activity. To evaluate possible underlying proteases, six amino acids before and after the semi-specific cleavage point were evaluated in relation to their natural abundance (Figure 6). In position P1 of the peptide, aliphatic (Valin) and aromatic (Tyrosine) amino acids occurred more frequently (up to 5 times more often). Up- and downstream of the cleavage point aliphatic amino acids overweight (Phenylaniline and Histidine). Therefore, the cleavage pattern observed is not dominated by activity of MMPs, but rather by proinflammatory proteases such as cathepsin G or neutrophil elastase which prefer aliphatic amino acids in the P1 position.

**Figure 5.**
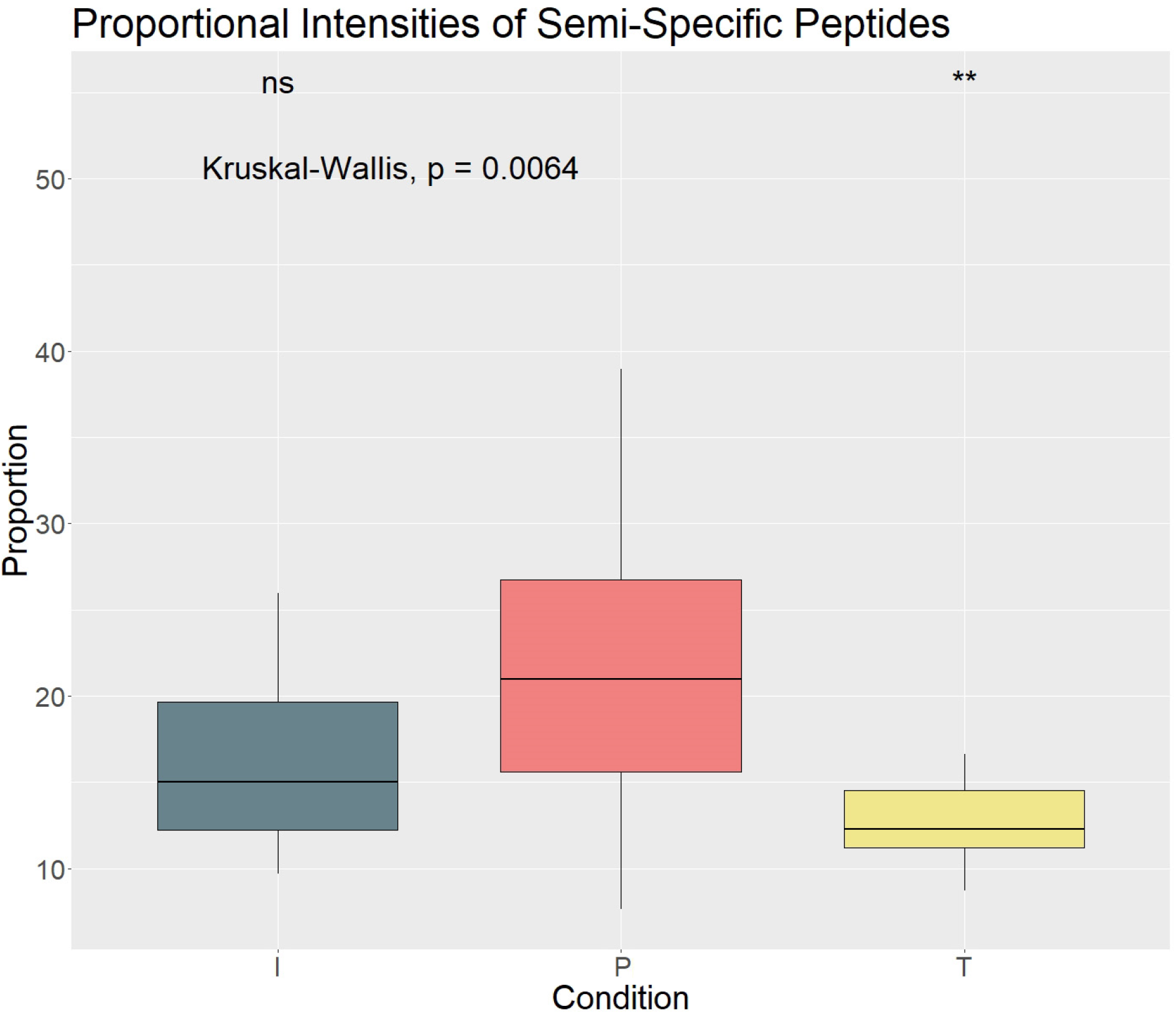
Proportional intensities of semi-specific peptides in different conditions. Kruskal-Wallis and subsequent Dunns post-hoc tests revealed significant differences between peri-implanitits (P) and healthy teeth (T) (p = 0.0057).

**Figure 6.**
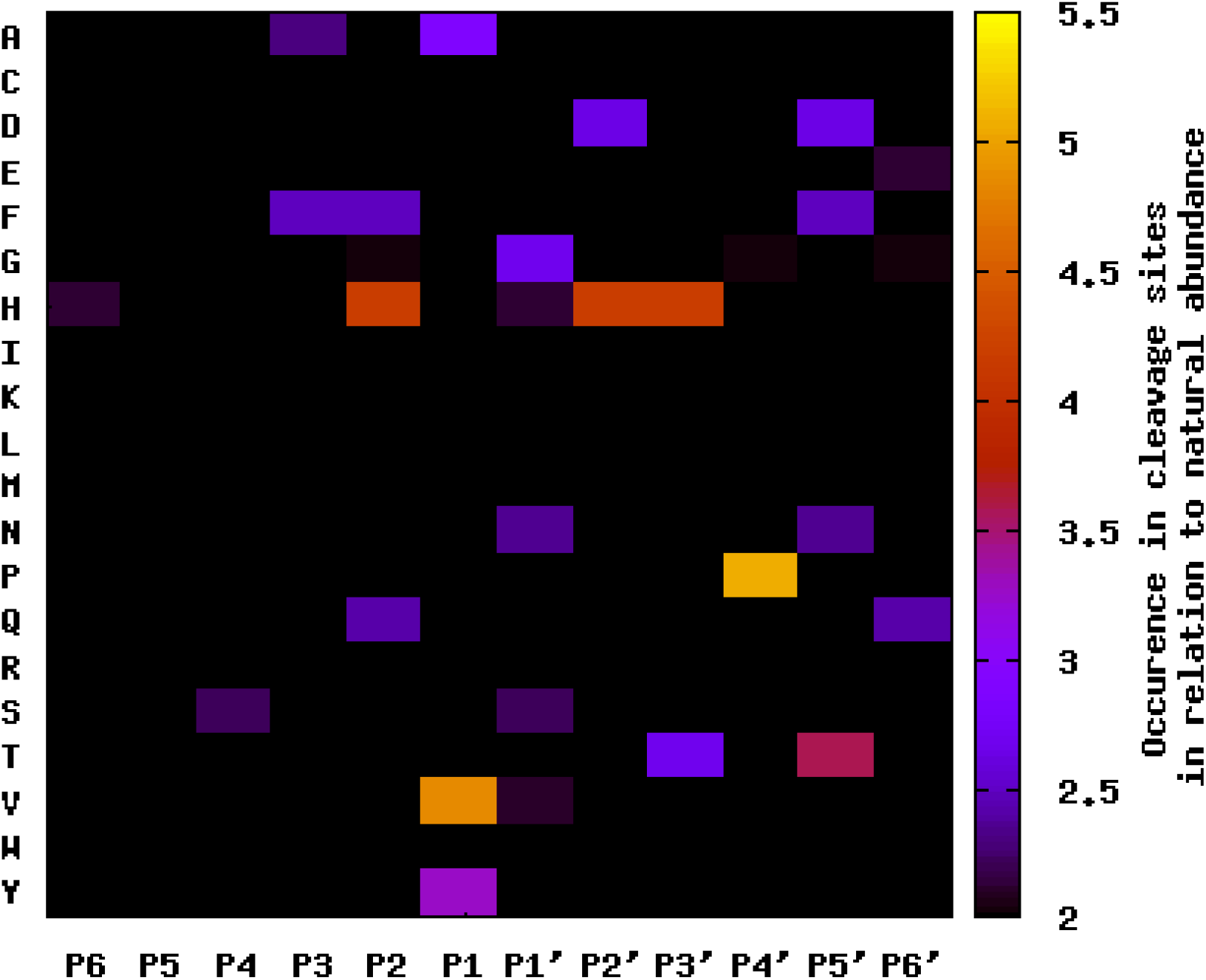
Heatmap of distribution of amino acids at different positions in relation to semi-specific cleavage points. *LIMMA* was performed to compare frequency of amino acids to natural abundance P_adj_ ≤ 0.05. X-axis represents amino acid positions P1-P6 up- and downstream of cleavage sites. Y-axis represents amino acids in one letter nomenclature. P1 shows increased occurrence of aromatic and aliphatic amino acids, such as valine (V) or tyrosine (Y).

### Bacterial Proteins

Further, we sought to investigate the presence of bacterial proteins in the crevicular fluid. A spectral library representing the human proteome supplemented with the proteome of the human oral cavity as defined by the NIH Human Microbiome Project Reference Genome Sequence Database was created^28^. This approach only serves as a proxy for the human oral microbiome, this approach does not capture species that are absent from these data sources. 334 bacterial proteins were identified and quantified. To evaluate differences in intensities of bacterial proteins, proportional intensities of all bacterial proteins found were calculated per sample (**Fehler! Verweisquelle konnte nicht gefunden werden**.1). Kruskal-Wallis rank test revealed significant differences between conditions (P, I and T) and Dunns post-hoc-test indicated significant differences between P and both healthy conditions (T and I). Furthermore, missingness of bacterial proteins was calculated. In the healthy conditions, missingness was especially high with nearly no proteins identified in all samples. Missingness was lower in the peri-implantitis subgroup, yet still more than 75% of all proteins were missing in more than half of all samples. Again, Kruskal-Wallis rank test and subsequent Dunns post-hoc-test showed significant differences between the diseased and both healthy conditions (Supplementary Figure 2).

## Discussion

The objective of the present study was to compare the P, I and T crevicular fluid on the proteome level. By applying MS-based proteomics, more than 2300 distinct proteins were identified, exceeding previous proteomic studies on crevicular fluid GCF ^23,29,30^ and revealing proteomic differences between healthy and diseased implants.

Data-independent acquisition was applied for quantitative proteomics of crevicular fluid in the present study. DIA has proven to produce more robust results, less vulnerable to stochastic biases, undersampling or limited reproducibility than conventional data-dependent acquisition (DDA) ^31–33^. The results of this study suggest that the DIA-approach in combination with the protein extraction protocol used in this study may enable deeper proteome decryption, especially when dealing with small sample volume as in crevicular fluid ^34,35^.

PLS-DA and *LIMMA* indicated little alterations in the proteomic composition of PICF/GCF of healthy teeth and healthy implants. In contrast, 59 proteins were identified as upregulated and 31 downregulated in peri-implantitis when compared to healthy implants. To date, no study has been published comparing the proteomic crevicular fluid composition of healthy and diseased implants, nor of healthy implants and healthy teeth. Esberg et al. found 42 proteins to be increased in case of implant loss after peri-implantitis therapy in comparison to peri-implantitis therapy success ^24^. Several studies have described altered protein expressions between healthy and periodontally diseased teeth and indicated a discrimination of periodontally healthy and diseased teeth on the proteome level ^22,35,36^.

In the present study, gene ontology analysis revealed biological processes related to local immune response like “humoral immune response”, “extracellular matrix organization” or “regulated exocytosis” to be upregulated in peri-implantitis. This is in accordance with the classic depiction of peri-implantitis as a local inflammation, orchestrated by the host immune response^37^. Interestingly, biological processes like “humoral immune response” and “defense response” were also found increased in a transcriptomic study on peri-implant soft tissue38. Despite the different experimental setups (proteomics vs. transcriptomics), this could suggest that the crevicular fluid mirrors the inflammatory state of the surrounding tissue, hence further emphasizing how PICF could help to monitor peri-implant health and disease. It is noteworthy that Becker et al. describe distinct differences in the biological processes observed in P and periodontitis, with the latter showing increased proliferative pathways38. It remains to future research to study these differences on the proteomic level and in the PICF.

Several proteins expressed by neutrophiles (myeloperoxidase, bacterial permeability-increasing protein, azurocidin, cathelicidin antimicrobial peptide), inheriting antibacterial activity, were identified in higher abundance in the peri-implantitis proteome. Azurodicin is expressed by neutrophiles and shows antimicrobial activity against Gram-negative species, including *P. aeruginosa* and *A. actinomycetemcomitans*, by binding lipopolysaccharides (LPS) ^39,40^. Likewise, myeloperoxidase is expressed in neutrophiles and acts synergistically with azurocidin ^41^. In periodontology, azurodicin and myeloperoxidase were found to be elevated in GCF of periodontitis sites, using both ELISA and MS-based proteomics ^30,42,43^. Also, myeloperoxidase was found more abundant in PICF of implants with peri-implantitis ^44^, which is in accordance with the findings of the present study.

Higher proteolytic activity was found in peri-implantitis, with cleavage patterns pointing to typical inflammatory proteases cathepsin G and neutrophil elastase. In inflammatory diseases, endogenous proteolysis inherits an important role, which has not been assessed for peri-implantitis previously. Both cathepsin G and neutrophile elastase are secreted by neutrophiles in azurophilic granules alongside neutrophilic proteins ^45^. Neutrophile granulocytes have been found in peri-implantitis lesions both in higher proportion and in higher total amounts when compared to periodontitis ^18,46^. Increased proteolytic activity by cathepsin G and/or neutrophil elastase as well as upregulation of azurocidin and myeloperoxidase, as presented here, underlines elevated neutrophile activity in peri-implant inflammation.

In this study, proteins involved in inflammatory processes were detected to be upregulated e.g. S100P, leukotriene A(4) hydrolase (LTA4H) or lymphocyte specific protein 1 (LSP-1), which have not been described in crevicular fluid before ^24,30,47^. LSP-1 and LTA4H inherit proinflammatory properties through their contribution to migration and lamellipodia formation of macrophages ^48,49^ and conversion of leukotriene A4 into proinflammatory leukotriene B4 ^50^, respectively. S100P is a calcium binding protein localized in the cytoplasm and the nucleus. Calcium-binding of S100P leads to activation of several target proteins, such as RAGE (receptor for advanced glycation end points) ^51^. S100P plays a central role in inflammation and bone homeostasis and may resemble a link between immune response and crestal bone loss ^52^. Since this is the first study to find elevated levels of S100P in case of P, further investigations are necessary to evaluate its role in peri-implantitis.

Interestingly, other members of the S100-family (S100A6, S100A7, S100A8, S100A9) have frequently been found upregulated in GCF of periodontally diseased teeth ^30^. Besides S100P, no significantly elevated levels of calcium-binding S100-proteins were detected. In contrast S100A11 was significantly decreased in P whereas in periodontitis S100A11 was found elevated in GCF ^53^. These differences regarding S100-proteins may indicate distinct differences of the crevicular fluid composition between periodontitis and peri-implantitis on the proteome level.

Matrix metalloproteinases (MMPs) 8 and 9 were identified in all samples and most abundant in the diseased subgroup (P), yet without significance. MMPs have frequently been observed to be elevated at periodontally diseased sites, with some publications also describing upregulation in case of peri-implantitis ^54–58^. Previous studies mostly focused on MMP8, a collagenase, secreted by neutrophils ^57–59^. No MMP dominated fingerprint of endogenous proteolysis was detected in the present study. Since rapid degradation of MMP-cleavage products *in vivo* cannot be ruled out, this finding does not allow to rule out increased MMP activity in peri-implantitis lesions. Activity assays of MMPs could be more suited to evaluate the role of MMPs in tissue degradation of peri-implantitis.

In this study, mass spectrometry-based assessment of bacterial proteins in PICF was performed for the first time, demonstrating that the relative intensity of bacterial proteins was increased in peri-implantitis. Elevated proportions of bacterial proteins were also found by Bostanci et al. ^22^ for periodontitis. The proportion of samples a given bacterial protein was not identified in (missingness) was high in all conditions. Almost no bacterial proteins were identified in more than 50% of the samples of I and T, indicating an inhomogeneous distribution of bacterial proteins. This is in accordance with microbiome studies, pointing to a diverse and ambiguous microbiomes associated with peri-implantitis. Therefore, it is debatable whether bacterial proteins may serve as diagnostic indicators of P ^60,61^.

The method presented may not only enable the identification of biomarkers for early detection of P, yet proteomic data could also help to stratify between patients and help in peri-implantitis risk assessment. The possibility to assess proteomic clusters with the help of MS could help to derive patient-specific risk prediction models. In a recent review, Steigmann et al. describe the possibilities given by advances in biotechnology and artificial intelligence to make use of biomarker information for clinical monitoring of oral diseases^62^. A comprehensive assessment of the PICF proteome, as demonstrated in the present article for the first time, may be an important step towards a patient specific precision medicine regarding peri-implant diseases. The GCF composition of periodontitis affected teeth has already been assessed by several studies, yet in the present study only healthy teeth were included. A direct comparison of diseased implants and teeth could lead to new insights into the disease’s biology, similarities and differences, and should therefore be subject of future research.

This study was designed as a proof of concept to demonstrate the potency of mass spectrometry-based proteomics on crevicular fluid of peri-implantitis. Therefore, it is limited by its small cohort size and statistical results should be considered carefully. Furthermore, crevicular fluid volume was not assessed before protein extraction and absolute protein concentrations in the crevicular fluid can’t be derived from this analysis.

Proinflammatory cytokines, as discussed by a variety of publications using antibody-based protein detection in periodontitis and peri-implantitis, were not identified in the present study. Previous MS-based studies on periodontitis similarly did not quantify these proteins^63–65^. Missingness of cytokines in MS-based proteomic studies has frequently been attributed to low concentrations of the molecules of interest ^22,64,65^. Previous studies applying targeted methods successfully demonstrated the role of proinflammatory cytokines in peri-implant and periodontal disease, however these proteins could not be used for diagnostic purposes due to low specificity and sensitivity ^57,66,67^.The method presented here enables to gain a broader view of the PICF proteome and may therefore be more suitable for detection of potential biomarkers. A combination of MS-based analyses and classical assaying for inflammatory proteins could enable an even deeper decryption of the proteomic composition. Furthermore, targeted MS-based approaches could be applied for cytokine identification.

Data presented here shall aid for power calculations of future investigations. Larger cohorts are needed to correlate proteomic data with patient individual clinical data to enable evaluations of potential etiological pathways. Patient individual differences related to age, sex, preexisting diseases or history of smoking have not been considered for the analysis. However, the influence of patient specific systemic factors on the GCF or PICF composition has not been evaluated adequately in literature. Proteomic findings of this study are limited to the composition of PICF and GCF. More research seems to be necessary to evaluate the connection between the PICF proteome and inflammatory tissue composition ^68^.

## Conclusion

The PICF proteome of peri-implantitis affected sites exhibits an inflammatory fingerprint when compared to healthy implants and teeth. Healthy implants and teeth share strong similarities in their proteome. The method demonstrated may pave the way to a more profound understanding of disease biology, new diagnostic and prognostic tools and a patient-specific therapy of peri-implantitis in the future.

## Acknowledgement

OS acknowledges funding by the German Research Foundation (Deutsche Forschungsgemeinschaft (DFG), Bonn, Germany)(SCHI 871/17-1 (Heisenberg program); SCHI 871/15-1 (individual research grant program); GR 4553/5-1 (individual research grant program); PA 2807/3-1 (individual research grant program); NY 90/6-1 (individual research grant program); INST 39/1244-1 (P12) (program: collaborative research centers); INST 39/766-3 (Z1) (program: collaborative research centers); 423813989/GRK2606 “ProtPath” (program: research training groups); Project-ID 441891347-SFB-1479; Project-ID 431984000 – SFB 1453) (program: collaborative research centers) mat; the ERA PerMed program (BMBF, 01KU1916, 01KU1915A); the German-Israel Foundation (grant no. 1444); the German Consortium for Translational Cancer Research (project Impro-Rec); and the Fördergesellschaft Forschung Tumorbiologie (promotional society for tumor biology research, Freiburg, Germany) (projects ILBIG and NACT).

## Authors’ Contributions

TH: Proteomic analysis, Statistics, Writing.

KN: Conceptualization and Idea, Surgeries, Administration, Review and Editing.

GI: Review and Editing.

OS: Proteomic analysis, Statistics, Review and Editing.

TF: Conceptualization and Idea, Surgeries, Administration, Writing.

## Footnotes

** Periopaper, Oraflow Inc., Hewlett, NY, USA

† Lysyl Endopeptidase ®, Mass spectrometry grade, Fujifilm/WAKO

‡ Mass spectrometry grade, Promega

§ Preomics

** Q-Exactive Plus Mass spectrometer, Thermo Scientific, San Jose, CA, US

†† Thermo Scientific, San Jose, CA, US

‡‡ Biognosys, Schlieren, Switzerland

§§ Dr. Maisch HPLC GmbH, Ammerbusch, Germany

*** Biognosis, Schlieren, Switzerland

