## Supplementary Figure 1 for "The Impact of Peri-implantitis on the Proteome Biology of Crevicular Fluid: A pilot study"

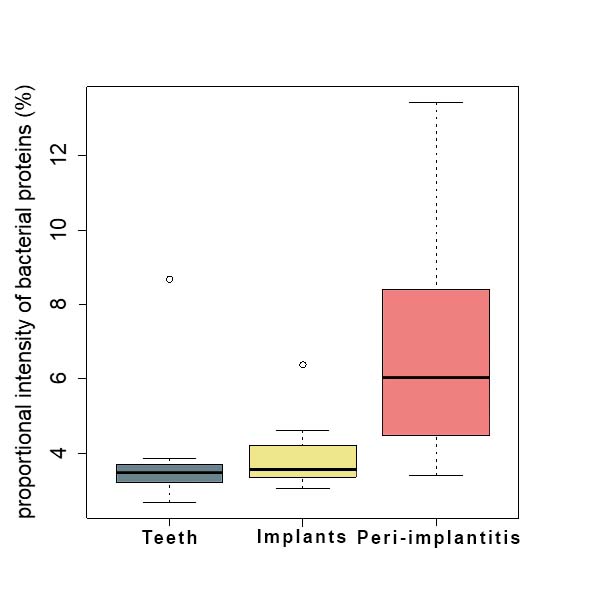

*****

*****

**Supplementary Figure 1** - Proportional intensities of bacterial proteins in relation to all proteins quantified (bacterial and human). Kruskal-Wallis rank test and Dunns post-hoc test indicate significant differences between PI and healthy subgroups. Peri-implanitits (PI), healthy teeth (T) and healthy implants (I).
