## Supplementary Figure 2 for "The Impact of Peri-implantitis on the Proteome Biology of Crevicular Fluid: A pilot study"

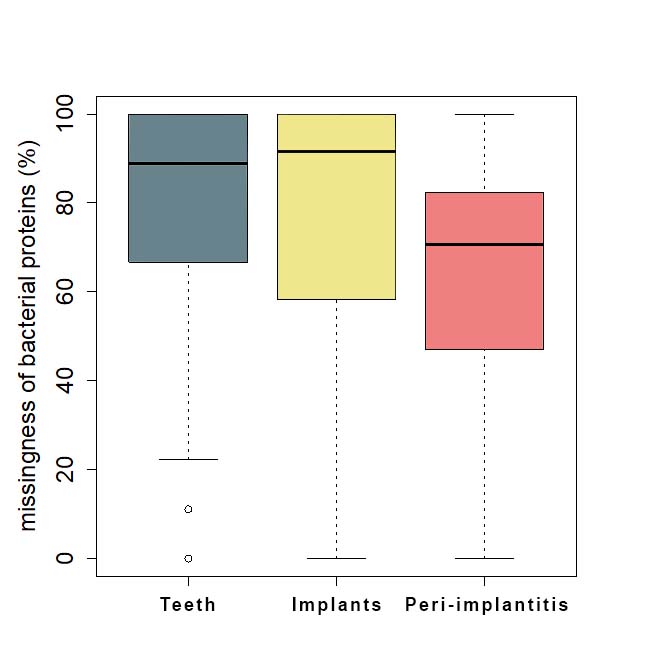

*****

*****

**Supplement Figure 2 -** Missingness of bacterial proteins. Especially high missingness in healthy subgroups. Kruskal-Wallis and Dunns post-hoc indicate lower missingness in the P subgroup. Peri-implanitits (P), healthy teeth (T) and healthy implants (I).
