## Supplementary Table 1 for "The Impact of Peri-implantitis on the Proteome Biology of Crevicular Fluid: A pilot study"

**Supplementary Table 1** – Patient data. 14 Patients with 38 samples were included in the study.

| Patient-Number | Sex | Age | P  localization | Implant localization | Tooth localization | Sample Number | Nicotine | Radiatio  Head or Neck | Pre-existing diseases | Current Medication | History of Pharmaceutical Cancer Treatment |
| --- | --- | --- | --- | --- | --- | --- | --- | --- | --- | --- | --- |
| PP3 | W | 74 | 19 (36) |  |  | P_03 | Y (15PY) | Y (60 Gy; 1989) | State after Breast Cancer (2011)  State after Adenoid Cystic Carcinoma (1989) | N | Y (Tamoxifen 2011 – 2016) |
|  |  |  |  | 10 (22) |  | I_03 |  |  |  |  |  |
|  |  |  |  |  | 27 (43) | T_03 |  |  |  |  |  |
| PP5 | W | 66 | 3 (16) |  |  | P_05 | N | N | N | N | N |
|  |  |  |  | 28 (44) |  | I_05_a |  |  |  |  |  |
|  |  |  |  | 9 (21) |  | I_05_b |  |  |  |  |  |
|  |  |  |  |  | 8 (11) | T_05 |  |  |  |  |  |
| PP6 | M | 70 | 3 (16) |  |  | P_06 | N | N | Factor V-Leiden | Macumar | N |
|  |  |  |  | 5 (14) |  | I_06 |  |  |  |  |  |
|  |  |  |  |  | 7 (12) | T_06 |  |  |  |  |  |
| PP7 | M | 77 | 3 (16) |  |  | P_07 | N | N | State after Apoplex,  State after Prostate Cancer (2003) | Clopidogrel, Ramipril | N |
|  |  |  |  | 30 (46) |  | I_07 |  |  |  |  |  |
|  |  |  |  |  | 5 (14) | T_07 |  |  |  |  |  |
| PP9 | W | 68 | 6 (13) |  |  | P_09 | Y (25PY) | N | N | N | N |
|  |  |  |  | 24 (31) |  | I_09 |  |  |  |  |  |
|  |  |  |  |  | 11 (23) | T_09 |  |  |  |  |  |
| PP13 | M | 74 | 15 (27) |  |  | P_13 | N | N | Hypertension | N | N |
|  |  |  |  | 12 (24) |  | I_13 |  |  |  |  |  |
|  |  |  |  |  | 8 (11) | T_13 |  |  |  |  |  |
| PP8 | W | 80 | 12 (24) |  |  | P_08 | N | N | Hypertension, Cataract,  Hypertyroidism | L-Thyroxin, Bisoporolol | N |
|  |  |  |  | 13 (25) |  | I_08_a |  |  |  |  |  |
|  |  |  |  | 14 (26) |  | I_08_b |  |  |  |  |  |
|  |  |  |  | 15 (27) |  | I_08_c |  |  |  |  |  |
| PP14 | M | 70 | 6 (13) |  |  | P_14 | N | N | N | N | N |
|  |  |  |  | 4 (15) |  | I_14 |  |  |  |  |  |
| PP15 | W | 85 | 6 (13) |  |  | P_15 | N | N | N | N | N |
|  |  |  |  | 11 (23) |  | I_15 |  |  |  |  |  |
| PP10 | M | 52 | 9 (21) |  |  | P_10 | Y  (18 PY) | N | N | N | N |
|  |  |  |  |  | 7 (12) | T_10 |  |  |  |  |  |
| PP12 | M | 77 | 31 (47) |  |  | P_12_a | N | N | N | Padraxa | N |
|  |  |  | 29 (45) |  |  | P_12_b |  |  |  |  |  |
|  |  |  |  |  | 25 (41) | T_12 |  |  |  |  |  |
| PP16 | M | 60 | 30 (46) |  |  | P_16 | N | Y (75 Gy; 2009) | State after Tongue Carcinoma (2008) | Pantoprazol, L-Thyroxin | Y (Cis-Platin 03-04/2009) |
|  |  |  |  |  | 7 (12) | T_16 |  |  |  |  |  |
| PP1 | M | 70 | 8 (11) |  |  | P_01_a | N | N | N | N | N |
|  |  |  | 3 (16) |  |  | P_01_b |  |  |  |  |  |
|  |  |  | 30 (46) |  |  | P_01_c |  |  |  |  |  |
| PP11 | W | 66 | 30 (46) |  |  | P_11 | N | N | N | N | N |

Abbreviations: W = female; M = male; N = No (does not apply); Y = Yes (does apply); PY = Pack Years, Gy = Gray. Age at date of sampling. Localization of teeth and implants according to ADA notation; FDI-notation in brackets.
