## Supplementary Table 2 for "The Impact of Peri-implantitis on the Proteome Biology of Crevicular Fluid: A pilot study"

| **Protease elevated in periimplantits** | **p_mod_** | **p_Adj_** |
| --- | --- | --- |
| Leukotriene A-4 hydrolase (LTA4H) | 0.00 | 0.04 |
| Gamma-glutamyl hydrolase | 0.00 | 0.05 |
| Neutrophil collagenase (Matrix metalloproteinase-8, MMP-8) | 0.04 | 0.14 |
| Matrix metalloproteinase-9 (MMP-9) | 0.04 | 0.15 |
| Acylamino-acid-releasing enzyme (AARE) | 0.06 | 0.18 |
| Neutrophil elastase (Human leukocyte elastase) | 0.27 | 0.46 |
| Dipeptidyl peptidase 1 (Cathepsin C) | 0.73 | 0.83 |
| Dipeptidyl peptidase 3 | 0.93 | 0.95 |

**Supplementary Table 2** - LIMMA of Proteases identified in peri-implantitits. 8 proteins with proteolytic activity were identified as upregulated in peri-implantitis. FDR-correction reduced on proteolytic enzymes (padj).
