## Supplementary Table 3 for "The Impact of Peri-implantitis on the Proteome Biology of Crevicular Fluid: A pilot study"

**Supplementary Table 3** – Enriched proteins in peri-implantitis when compared to healthy implants based on LIMMA.

| Uniprot ID | P_adj_ | Fold enriched in P |
| --- | --- | --- |
| A0A0C4DH43 | 0.04295996 | 2.65718203 |
| O15511 | 0.04881405 | 1.77904035 |
| O75695 | 0.04312712 | 2.61761471 |
| O75923 | 0.04036429 | 2.36560971 |
| O75955 | 0.04295996 | 2.45845219 |
| O95445 | 0.03307215 | 2.17979049 |
| P00492 | 0.04312712 | 1.79230743 |
| P00918 | 0.04312712 | 5.1437266 |
| P02042 | 0.04881405 | 8.56382365 |
| P02750 | 0.04820616 | 2.13942235 |
| P02753 | 0.04881405 | 2.33649237 |
| P04040 | 0.04881405 | 2.88468336 |
| P05107 | 0.04036429 | 2.4741577 |
| P05155 | 0.03307215 | 2.1633903 |
| P05164 | 0.04102584 | 2.61163561 |
| P05546 | 0.04312712 | 3.0876966 |
| P06737 | 0.04102584 | 2.00390883 |
| P06744 | 0.04623636 | 1.67691079 |
| P07738 | 0.04471604 | 6.61277911 |
| P08133 | 0.04312712 | 2.63180594 |
| P10153 | 0.03646101 | 2.14550202 |
| P11215 | 0.00559858 | 3.1096591 |
| P11413 | 0.031483 | 1.92142324 |
| P12429 | 0.02380754 | 1.87940878 |
| P13716 | 0.04572494 | 2.40448728 |
| P16083 | 0.04312712 | 2.70650776 |
| P17213 | 0.03673849 | 2.67062977 |
| P18084 | 0.01516677 | 3.57565621 |
| P20160 | 0.02334487 | 2.43590022 |
| Q92820 | 0.04623636 | 2.25711505 |
| P09960 | 0.04312712 | 1.55074406 |
| P25815 | 0.00694557 | 3.28141451 |
| P26583 | 0.04300998 | 2.00721447 |
| P27105 | 0.04881405 | 1.78312496 |
| P28676 | 0.03646101 | 2.41412059 |
| P29401 | 0.04312712 | 1.91743907 |
| P30046 | 0.04036429 | 1.61138547 |
| P30048 | 0.04312712 | 1.64295144 |
| P30626 | 0.04102584 | 1.41127563 |
| P31146 | 0.02334487 | 2.82425522 |
| P33241 | 0.04312712 | 2.23694054 |
| P36222 | 0.04312712 | 2.22084444 |
| P37837 | 0.01568107 | 1.60394784 |
| P41218 | 0.04036429 | 1.78192011 |
| P49913 | 0.01568107 | 2.356056 |
| P52566 | 0.02334487 | 1.83683045 |
| P52790 | 0.04295996 | 2.35527628 |
| P69905 | 0.04036429 | 9.21326452 |
| P80723 | 0.01568107 | 2.48547012 |
| Q10588 | 0.04312712 | 2.48571664 |
| Q13228 | 0.04312712 | 3.63643217 |
| Q14254 | 0.02380754 | 2.32787441 |
| Q6P4A8 | 0.02894062 | 2.74515092 |
| Q99439 | 0.04036429 | 1.68384081 |
| P67812 | 0.03928548 | 2.01356665 |
| Q9H4G4 | 0.04102584 | 2.04635567 |
| Q9HD89 | 0.03307215 | 3.16582353 |
| Q9HDC9 | 0.00694557 | 2.88406407 |
| Q9Y2J8 | 0.04295996 | 2.15058364 |

Protein names as Uniprot identifiers. P_adj_ according to LIMMA.
