## Supplementary Table 4 for "The Impact of Peri-implantitis on the Proteome Biology of Crevicular Fluid: A pilot study"

**Supplementary Table 4** – Downregulated proteins in peri-implantitis when compared to healthy implants based on LIMMA.

| Uniprot ID | P_adj_ | fold decreased in P |
| --- | --- | --- |
| O14579 | 0.04312712 | 0.44808162 |
| O43776 | 0.04312712 | 0.50067601 |
| O75964 | 0.031483 | 0.50949977 |
| O95834 | 0.04295996 | 0.62573453 |
| P04439 | 0.04471604 | 0.42410331 |
| P07339 | 0.00559858 | 0.56523966 |
| P07355 | 0.00694557 | 0.45854447 |
| P04632 | 0.04406981 | 0.65965342 |
| P07858 | 0.01568107 | 0.43462133 |
| P07900 | 0.04312712 | 0.64436532 |
| P09758 | 0.04623636 | 0.53644859 |
| P10599 | 0.04036429 | 0.56898967 |
| P13073 | 0.01235459 | 0.54592724 |
| P13797 | 0.04471604 | 0.44418262 |
| P15311 | 0.04572494 | 0.47860974 |
| P16152 | 0.04036429 | 0.504496 |
| P18124 | 0.04312712 | 0.61309693 |
| P30086 | 0.02334487 | 0.58899923 |
| P31949 | 0.04312712 | 0.57991978 |
| P40925 | 0.04036429 | 0.55676969 |
| P49189 | 0.04312712 | 0.435668 |
| P54920 | 0.04623636 | 0.69372621 |
| Q00325 | 0.04312712 | 0.61223321 |
| Q13200 | 0.04036429 | 0.68110328 |
| Q15149 | 0.03307215 | 0.53862439 |
| Q3LXA3 | 0.04312712 | 0.50043995 |
| Q99733 | 0.00310314 | 0.30715271 |
| Q99798 | 0.04624073 | 0.57988882 |
| Q9UHY7 | 0.031483 | 0.51674439 |
| Q9UK76 | 0.03646101 | 0.45284605 |
| Q9UL46 | 0.04036429 | 0.40555514 |

Protein names as Uniprot identifiers. P_adj_ according to LIMMA.
